# Circadian phase alignment promotes survival under predictive oxidative stress

**DOI:** 10.64898/2026.10.11.758360

**Authors:** Alexandra Neitz, Arnaud Taton, Jimmy Yu, Veronica Ghaly, Susan S. Golden

**Affiliations:** Center for Circadian Biology and Medicine, University of California, San Diego, La Jolla, CA 92093; Department of Molecular Biology, University of California, San Diego, La Jolla, CA 92093

## Abstract

Circadian clocks are endogenous time-keeping mechanisms that continue to oscillate with a period that approximates 24 h in the absence of cyclic environmental cues. This free-running period (FRP) varies from species to species and can vary within species across latitudinal clines. Because natural environments provide daily cues that reset circadian clocks to exactly 24 h, the importance of the FRP length is unclear. Using the cyanobacterium *Synechococcus elongatus* we show that the FRP length alters the alignment of a biological process with an environmental reference, known as phase angle of entrainment, across seasonal photoperiods, thereby repositioning stress-response programs relative to the external environment. This phase relationship affects the overall fitness of the organism: strains that position stress-response programs to peak at the time at which the stressor peaks outcompete their early- or late-peaking counterparts. Our results suggest that the FRP length is selected for its ability to adjust the phase angle of entrainment across seasonal photoperiods: the wild-type FRP length was uniquely plastic in its ability to adjust stress-response programs to coincide with the stressor in all photoperiod conditions.

**Significance Statement:** Circadian clocks are nearly universal biological timing systems that allow organisms to anticipate daily environmental changes, optimize environmental resources, and avoid environmental threats. We investigated which features of clock function promote survival fitness and which environmental conditions provide selection. A long-standing hypothesis is that the phase relationship between the internal timing mechanism and the external environment is critical for the circadian clock to confer a fitness advantage. However, because this phase relationship is influenced by the circadian period, tests of this hypothesis confound the two features. We experimentally decoupled the free-running period and the environmental phase relationship to show that the phase relationship, rather than the free-running period, confers fitness under environmental conditions that impose a selection.

## Introduction

Most organisms have internal biological clocks that allow them to anticipate daily changes in the environment caused by the rotation of the earth. The rhythmic manifestations of the activity of this clock are called circadian rhythms. A key feature of the circadian clock is its ability to ensure that physiological processes occur at the relevant time relative to the external rhythmic environment— a relationship called the phase angle of entrainment. Another feature of the circadian clock is that it continues to tick in the absence of environmental signals (i.e. is self-sustained) and under these “constant conditions”, the period of the clock differs slightly from 24 h and is called the free-running period (FRP). That circadian clocks confer fitness has been demonstrated, but the importance of the self-sustained property is more elusive because organisms in the wild rarely—if ever—encounter true constant conditions. Nonetheless, the self-sustained property is characteristic of circadian clocks across all kingdoms of life. The self-sustained feature may confer a selective advantage through the ability to match the cycle to 24 h (the process of entrainment) and predict cyclic environmental cues (1, 2). In order for the circadian clock to be able to align and entrain to the 24 h environment, the length of the FRP must be close to 24 h (3) and slight deviations of the FRP from 24 h may select for the maintenance of resetting mechanisms. According to the non-parametric theory of entrainment, when the FRP differs slightly from 24 h, the clock compensates with daily phase shifts, and these shifts determine the stable relationship of each process with a characteristic environmental reference. In other words, changes in the FRP alter the phase angle of entrainment.

The genetically tractable cyanobacterium *Synechococcus elongatus* PCC 7942 has been instrumental as a model organism in answering fundamental questions about the evolution and fitness of circadian clocks (3-7). Its molecular clock has been extensively studied and relies on a post-translational mechanism consisting of the interaction of the KaiA, KaiB, and KaiC proteins (8). Using *S. elongatus*, the prediction that the FRP alters the phase angle of entrainment was previously demonstrated experimentally in a 24 h day as well as in environmental cycles with periods that differ from 24 h (called T cycles) (3).

Another important aspect of the self-sustained property may be that it enables the phase angle of entrainment to be modified as daily day/night conditions change, for example as photoperiods change throughout the year. Work in cyanobacteria has shown that a self-sustained circadian clock shifts its phase angle of entrainment to track midday across seasonal photoperiods (9), whereas a genetic model of an hourglass (not self-sustained) clock does not (6). This seasonal adjustment of phase angle may be influenced by the FRP. In fact, studies in diverse organisms have found within-species variations in FRP along latitudinal clines, where species at latitudes away from the equator experience greater annual variation in photoperiod (10-12). However, it is still not known how the FRP influences the phase angle of entrainment in different seasonal photoperiods or to what extent the phase angle of entrainment (over FRP) contributes to survival and fitness.

Here, we address whether, and if so how, the phase angle of entrainment contributes to the fitness and survival of cyanobacteria across seasonal photoperiods. First, we show that we can change the phase angle of entrainment of the *S. elongatus* clock by changing the FRP in all photoperiods. Next, we show that the phase angle of entrainment confers a fitness advantage only under high reactive oxygen species (ROS) conditions. Finally, we show that the fitness advantage of the wild-type (WT) FRP is driven by the preparedness to cope with ROS accumulation at dusk.

## Results

### FRP controls phase of dusk gene expression across seasonal photoperiods

Previous research has shown that the FRP affects the phase angle of entrainment in 12:12 hour light-dark (LD) and non-24 h T cycles (3). However, it is unknown how the FRP changes the phase angle of entrainment across seasonal photoperiods, when the time to complete one cycle remains 24 h while the amount of light available across the day changes. To address this gap, we used a bioluminescent reporter of the *S. elongatus* clock gene promoter P*kaiBC* as a proxy for dusk-phase gene expression across FRP mutants in three seasonal photoperiods: summer (16:8 LD), equinox (12:12 LD), and winter (8:16 LD). The period mutants were generated by making point mutations to effect substitutions in KaiC: short-period (SP) KaiC-A87V and long-period (LP) KaiC-P236S (13) (Table S1, S2). We made two WT control strains to match the antibiotic resistances of the FRP mutants: WT with nourseothricin resistance (WT Nt), and WT with gentamycin resistance (WT Gm) (Table S1, S2). We found that the WT strains adjusted the acrophase (i.e. peak) of P*kaiBC* gene expression to occur within 2 h of dusk in all photoperiods, while the long-period mutant had delayed gene expression in all photoperiods (Figure 1A-C). The short-period mutant had advanced gene expression relative to the WT strains in summer and equinox (Figure A-B), but interestingly, was able to adjust the acrophase to occur within one hour of dusk in winter (Figure 1C). The acrophase of P*kaiBC* expression relative to dusk is correlated with the period across all seasons (Figure 1 D-F). Namely, the shorter the period, the earlier the acrophase occurred. We next asked how this FRP-associated timing of phase relationship affects the fitness of cyanobacteria.

**Figure 1.**
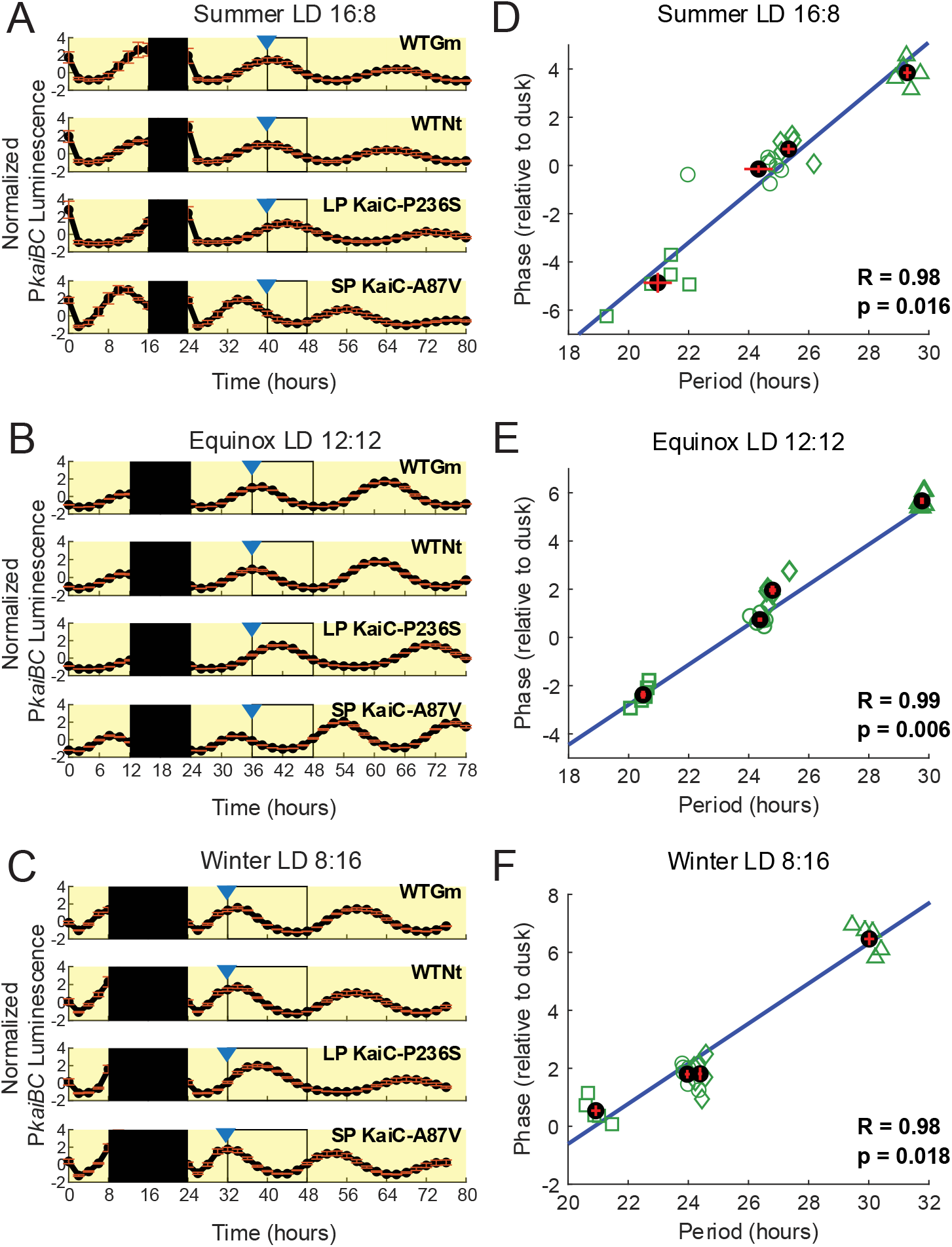
The FRP is associated with phase of expression from P*kaiBC* relative to dusk in all seasonal photoperiods. A-C.) Gene expression monitored from a bioluminescence reporter of P*kaiBC* in WT and FRP mutant strains in summer (A), equinox (B), and winter (C) photoperiods. The filled black box indicates the last night in an LD cycle. The black outline indicates first subjective night in constant light. The blue carrot indicates dusk. D-F.) The phase of expression from P*kaiBC* relative to dusk is correlated with the FRP in all seasons. Pearson’s correlation was used to calculate R and p-values for linear correlation (blue trendline). The green symbols represent individual replicates (○ WTNt, ◊ WTGm, □ SP KaiC-A87V, Δ LP KaiC-P236S). The black dots represent the mean of each strain, which are partially obscured by the red lines that represent the SEM. For summer and winter n_SP_=5, n_LP_=5, n_WTGm_=6, n_WTNt_=6, for equinox n_SP_=6, n_LP_=6, n_WTGm_=6, n_WTNt_=6.

### Under low-light conditions, FRPs that differ from WT have a fitness disadvantage in summer, but not in winter or equinox photoperiods

To test how the timing of clock gene expression affects fitness of cyanobacteria, we used growth-competition experiments of the WT against either the long-period or short-period mutants in three different photoperiods. In the summer photoperiod, WT outcompeted both the short-period and long-period mutants (Figure 2A). To see whether this difference in fitness can be explained by growth alone, we measured growth across time (Figure S1). Although the short-period mutant had a slight growth defect, the differences in growth among strains cannot explain the large difference in fitness that was observed in the growth-competition experiments. Previous reports have observed similar discrepancies between individual growth rates and growth-competition outcomes (3, 5).

**Figure 2.**
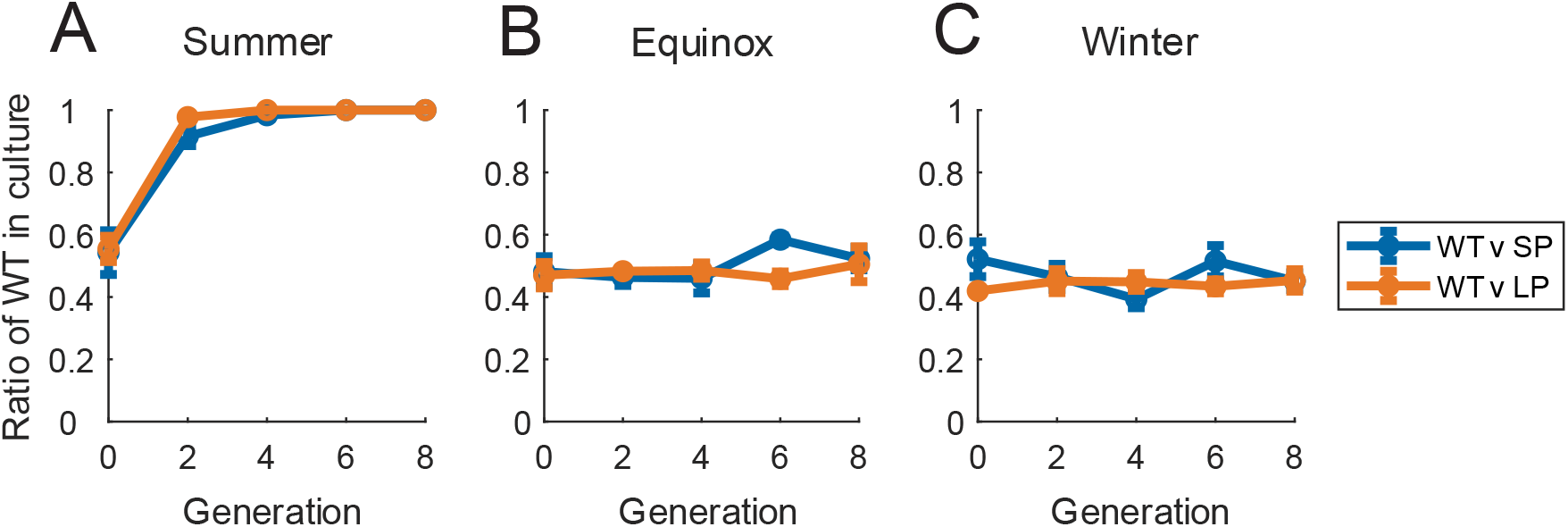
The WT FRP confers a fitness advantage in summer, but not equinox or winter photoperiods under low light conditions. Growth competition experiments between WT and long-period (LP) or short-period (SP) FRP mutants in A.) summer, B.) equinox and C.) winter photoperiods under low light (40 µmol/m^2^/s). Data represent mean±SEM, n=3 for all data points.

Surprisingly, we found that—despite differences in circadian gene expression between FRP strains—there was no advantage of FRP in equinox and winter photoperiods (Figure 2B, C). A critical role of the circadian clock in this organism is known to be the clearance of ROS at night (14). We reasoned that the difference in FRP advantage across photoperiods may be related to the accumulation of ROS across the day. Because ROS builds up during daytime photosynthesis, and the amount of time that cells are photosynthesizing across the day changes across seasonal photoperiods, we predicted that there are differences in ROS accumulation across photoperiods.

### Outcome of fitness experiments depends both on timing of dusk gene expression and differences in ROS accumulation at dusk

We hypothesized that ROS accumulates more in the summer than in the winter due to increased photosynthetic activity over the long days and that this difference in ROS explains the difference in fitness advantage of WT across seasons. We predicted that ROS can be decreased in the summer by increasing gas exchange and increased in the winter by increasing photosynthetic activity through increased light intensity. To test our predictions, we quantified ROS accumulation at dusk across 2 summer and 2 winter photoperiod conditions in WT and FRP mutant strains using the fluorescent marker H_2_DCFDA (Figure 3 A-D). To determine whether there are significant differences in accumulation of ROS across photoperiod conditions and strains, we ran a two-way ANOVA (Table S3). We found a significant difference among seasonal photoperiods (p=7.45e-15), but not across strains (p=0.26), and no significant interaction between photoperiod conditions and strain (p=0.10), indicating that the strains responded uniformly to the changing photoperiod. A Tukey’s post hoc comparison of seasonal photoperiod condition (Table S4) showed a significant difference between the amount of ROS accumulation between the summer (Figure 3A) and winter (Figure 3C) conditions that were used in Figure 2 (p=1.18e-11), suggesting that the WT FRP is only advantageous in stressful, high ROS conditions.

**Figure 3.**
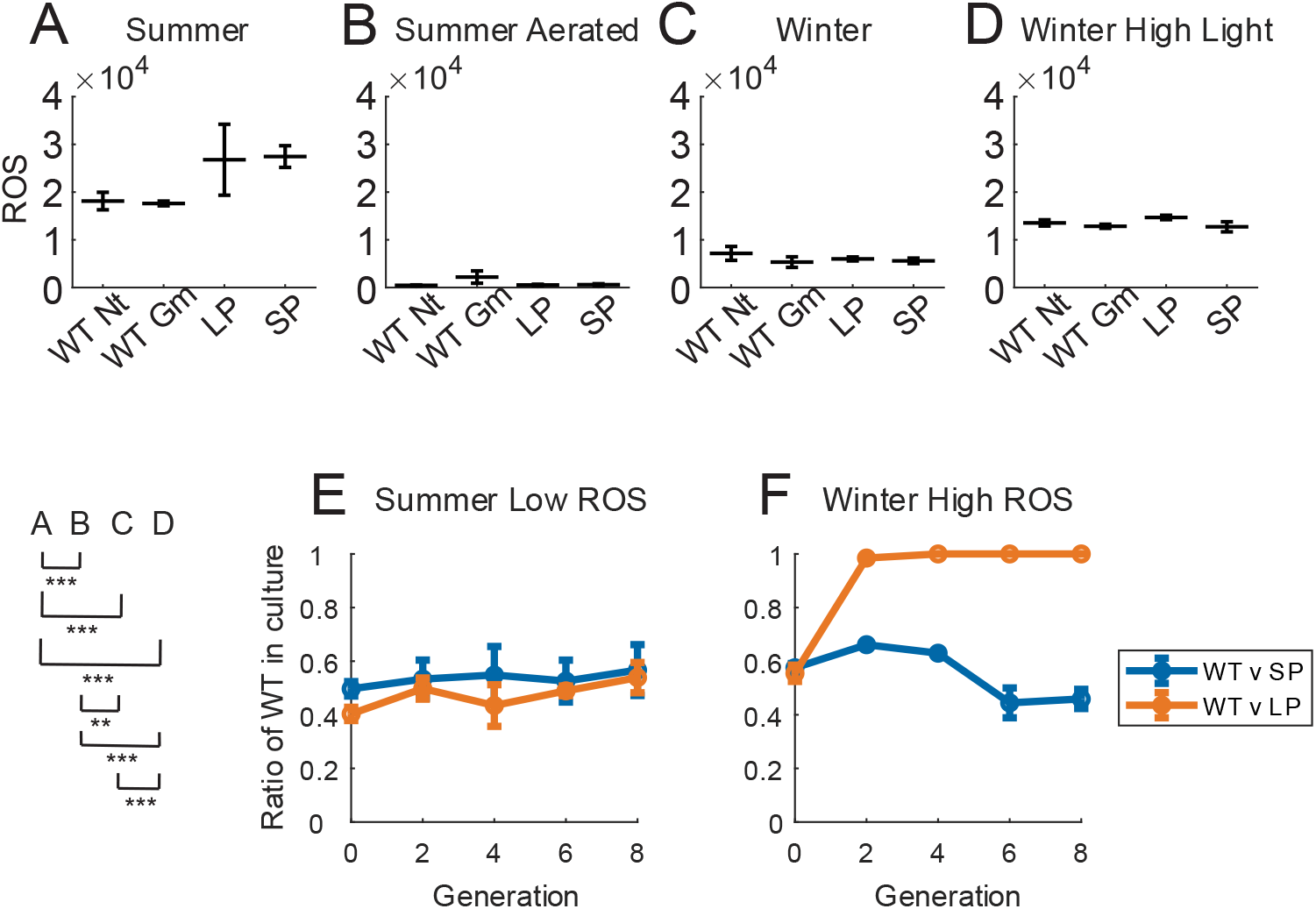
ROS accumulation drives fitness advantage of FRP length. A-D.) ROS accumulation at dusk across strains and experimental conditions (brackets on bottom left show p-values from two-way ANOVA followed by Tukey’s post-hoc test for comparisons between conditions, **p<0.01,***p<0.001). E-F.) Growth competition experiments between WT and long-period (LP) or short-period (SP) mutants in summer low ROS (aerated), and winter high ROS (high light). Data represent mean±SEM, n=3 for all data points.

We were able to significantly decrease ROS in the summer through aeration (p=8.42e-15, Figure 3A,B) and significantly increase ROS in the winter through increased light intensity (p=0.00012, Figure 3C,D). We hypothesized that increased accumulation of ROS drives the advantage of the WT FRP. We tested this hypothesis by performing growth-competition assays of the WT against either the long- or short-period mutants in the new summer and winter conditions. We found that lowered ROS in summer through aeration nullified the fitness advantage of WT (Figure 3E) and increased ROS in the winter drove the WT to outcompete the long-period strain (Figure 3F). Interestingly, WT did not outcompete the short-period strain in the winter high ROS condition (Figure 3F). We attribute this lack of advantage to the similarity of dusk gene expression phasing between the WT and short-period strains in the winter (Figure 1C). Together, our results indicate that the WT FRP is advantageous only in stressful, high-ROS conditions and that selection is acting on the phase angle of entrainment rather than the FRP.

### Ability to time ROS coping mechanisms explains the fitness advantage of the WT FRP

We next asked, how does the accumulation of ROS account for the advantage of the WT FRP? We did not detect a difference in the amount of ROS accumulation at dusk among strains (Figure 3A-D). Therefore, the amount of ROS accumulation at dusk alone does not explain the differences in fitness. Instead, we hypothesized that the timing of physiological processes that deal with ROS stress at night explain the differences in fitness between various FRP strains. The oxidative pentose phosphate pathway (OPPP) is critical for producing NADPH in the absence of photosynthesis, and flux through the OPPP is regulated by the circadian clock (14). The accumulation of fructose-6-phosphate (F6P) is an indicator of flux through the OPPP and the enzyme FBPase converts fructose-1,6-bisphosphate to F6P. The gene for the FBPase enzyme is *fbp* (Synpcc7942_2335), which is a dusk-peaking gene (15). The *fbp* gene resides within an operon that contains multiple OPPP-associated genes, making the operon’s regulation relevant to understanding the oxidative stress response. To assess the ability of the strains to cope with ROS at dusk, we monitored the timing of expression using a bioluminescence reporter driven by the native upstream operon promoter (P _2336_). *Synpcc_2336* has high dusk-peaking transcription (16) and its promoter contains a binding site for the master circadian transcription factor, RpaA (17). In the summer photoperiod, WT strains had an acrophase of P_2336_ expression occurring just before dusk (Figure 4A). Relative to WT, expression from P_*2336*_ was delayed in the long-period strain and advanced in the short-period strain (Figure 4A). Therefore, these results suggest that in summer the long-period strain is not prepared to cope with the accumulated ROS at darkness because there is a delay in the timing of turning on the OPPP. In contrast, the short-period mutant turns on the OPPP too early and by the time dusk arrives, the levels of expression from P_*2336*_ are already dropping.

**Figure 4.**
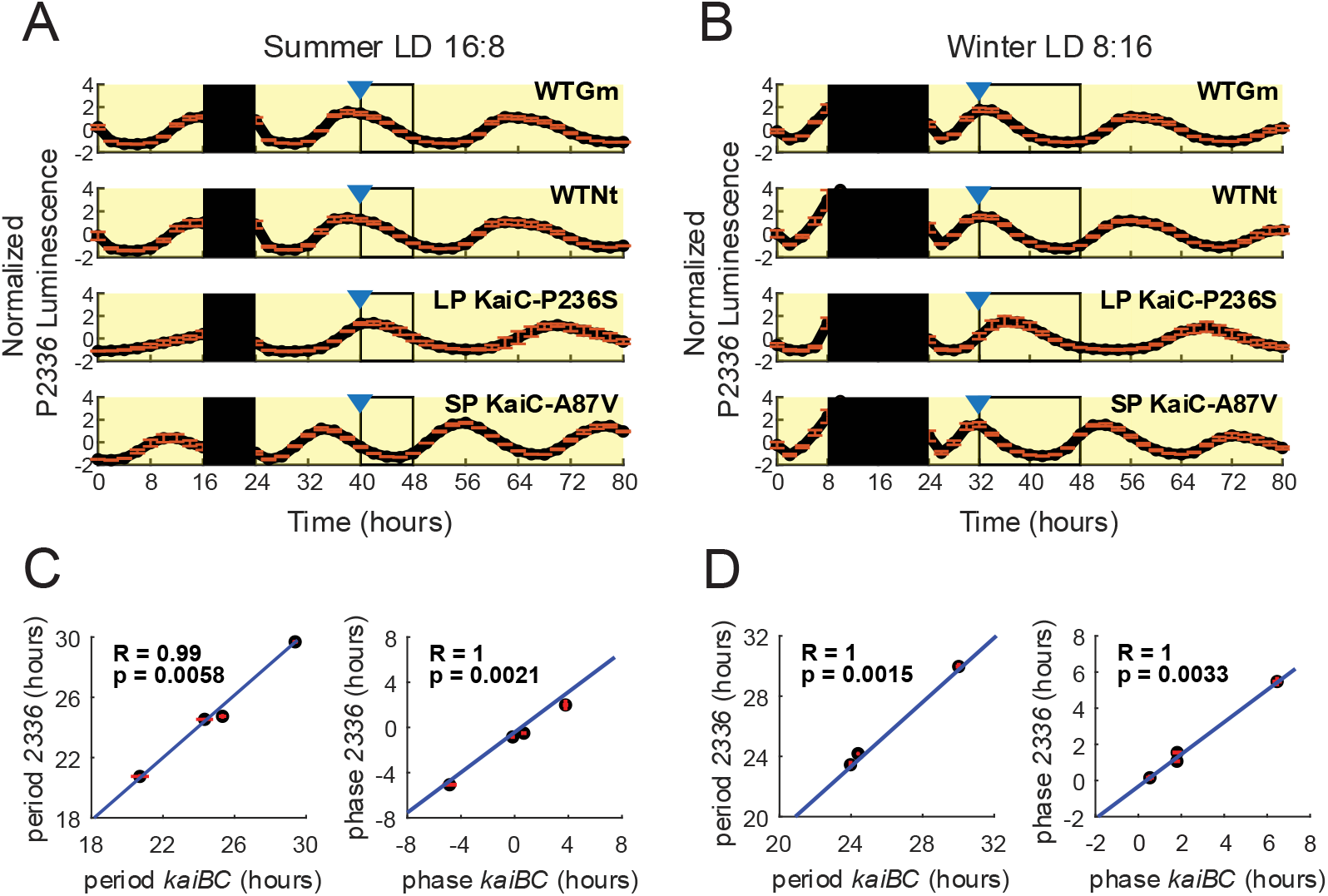
Gene expression from P_2336_, the promoter of a gene involved in redox balance at night, follows the circadian clock. A-B.) Gene expression monitored from a bioluminescence reporter of P_2336_ in WT and FRP mutant strains. The filled black box indicates the last night in an LD cycle. The black outline indicates first subjective night in constant light. The blue carrot indicates dusk. C-D.) Correlation between gene expression from P_2336_ and P*kaiBC*. Pearson’s correlation was used to calculate R and p-values for linear correlation (blue trendline). The black dots represent the mean of each strain, which are partially obscured by the red lines that represent the SEM. The P_2336_ summer sample sizes are as follows n_SP_=3, n_LP_=4,n_WTGm_=4, n_WTNt_=4. The winter P_2336_ sample sizes are as follows: n_SP_=5, n_LP_=4,n_WTGm_=6, n_WTNt_=7. The P*kaiBC* sample sizes for both winter and summer are: n_SP_=5, n_LP_=5, n_WTGm_=6, n_WTNt_=6.)

In winter, the acrophase of expression from P_*2336*_ was well aligned to dusk in the WT and short-period strains, but was several hours delayed relative to dusk in the long-period strain (Figure 4B). The similarity in the timing of expression in the WT and short-period strains may explain why there is no advantage of the WT over the short-period strain in the high ROS winter condition (Figure 3F). In contrast, the delay in expression from P_*2336*_ in the long-period strain indicates that it is not prepared to cope with ROS at dusk when the lights go off.

In WT cyanobacteria, different light environments can lead to differences in the timing of dusk gene expression (16). In other words, the same circadian genotype can have different dusk-peaking gene expression profiles depending on the external light environment. We therefore asked to what extent the timing of P_*2336*_ expression is controlled by the circadian clock in the winter and summer photoperiods. We found the period and phase of expression from P_*2336*_ were similar to those of P*kaiBC* in all strains in both winter and summer photoperiods, as expected for two direct targets of RpaA (Figure 4C-D).

If the fitness advantage of WT is driven by the ability of the strains to cope with ROS at dusk, then giving a dose of ROS at dusk should more negatively affect the survival of the mistimed strains than the properly aligned strains. We tested this prediction in a winter low-light photoperiod condition because the ROS levels are low. We added hydrogen peroxide to strains individually at dusk and quantified the ratio that survived (Figure 5A). A one-way ANOVA showed significant differences among strains (p=0.011, Table S5), and a Tukey’s post hoc test showed that the two WT strains, WT Nt and WT Gm, had significantly higher survival ratios than the long-period strain (p=0.018, p=0.041 respectively, Table S6). We did not detect a significant difference between the survival ratio of either of the WT strains and the short-period strain. We then asked whether the low survival of the long-period strain is due to an inherent inability of the strain to cope with ROS or the inability to cope with ROS at dusk. The long-period strain was exposed to hydrogen peroxide either at dusk or 4h after dusk, the time at which its acrophase of dusk-peaking gene expression occurs (Figure 5B). Its survival ratio was significantly higher when exposed 4 h after dusk than at dusk (student’s t-test of independent samples, t=-7.69, p = 9.17e-7). Notably, its survival to exposure 4 h after dusk was comparable to WT survival when WT cells were exposed at dusk. This pattern demonstrates that the long-period strain possesses the same underlying capacity to cope with ROS as the WT strain; however, because the long-period strain’s stress-response program is expressed at the wrong time, that capacity is only realized when the exposure happens to coincide with its mistimed peak.

**Figure 5.**
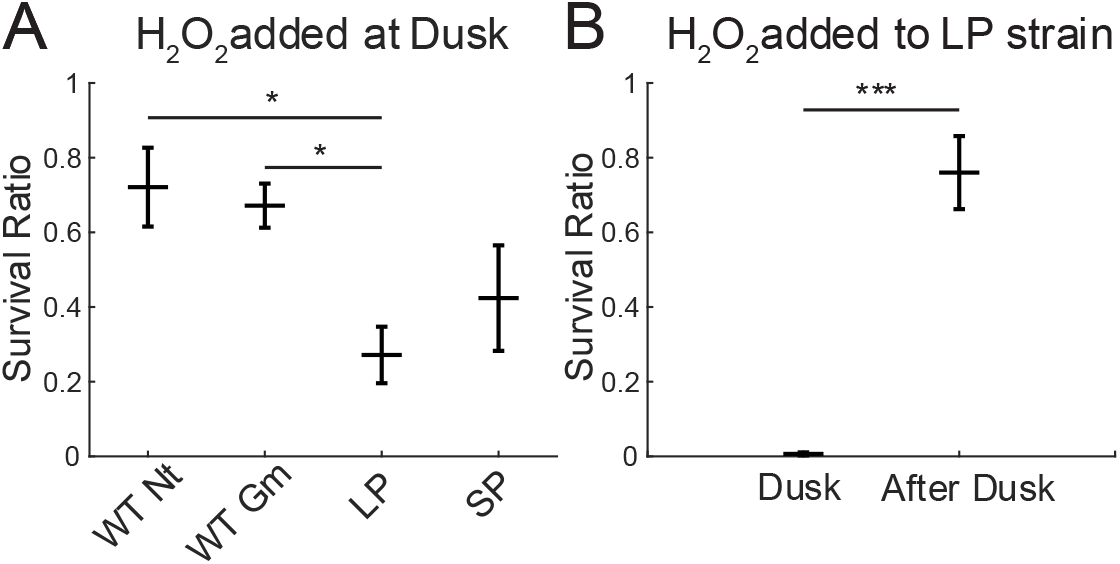
Survival to hydrogen peroxide exposure in a winter photoperiod. A.) Hydrogen peroxide exposure at dusk across all strains in a winter photoperiod. A one-way ANOVA followed by a Tukey’s post hoc test was used to determine p-values (*p<0.05). Data represent mean+SEM, n=8 for all data points. B.) Hydrogen peroxide exposure at dusk and 4 h after dusk in the long-period (LP) strain. A student’s t-test of independent samples was used to determine the p-value (***p<0.001). Data represent mean+SEM, n=9 for all data points.

Together these results highlight that proper temporal alignment of stress-response pathways is important for survival. Likely, the improper timing of expression of dusk-peaking genes, such as the OPPP pathway genes, in the long-period strain contributes to its fitness disadvantage in winter. On the other hand, the short-period strain is similar to WT in its timing of expression from P_*2336*_, which likely confers its ability to cope with ROS at dusk and explains why it does not have a fitness disadvantage vs WT in this condition.

## Discussion

The question of how circadian clocks promote fitness and how their interactions with the environment influence that advantage has been pursued since the discovery of intrinsic circadian timing (2). It has been shown previously that an FRP that matches that of the environment confers a fitness advantage (3). However, because the FRP has such a strong influence over the phase angle of entrainment it remained unanswered whether selection acts on the FRP per se, or the resulting phase relationship of processes with the environment. Here, the winter condition gave rise to the same phase angle of entrainment for two different FRPs, allowing us to identify the individual contributions of each. WT did not outcompete the short-period strain under the winter high ROS condition when the phase angle of entrainment between the two strains was similar. However, in the summer high ROS condition, when the phase angle of entrainment differed between the two strains, WT had an advantage over the short-period strain. Together these results strongly suggest that the selection acts not directly on the FRP but on the phase angle of entrainment. Furthermore, the flexibility of the phase angle of entrainment across seasonal conditions may have contributed to the selection of self-sustained oscillators. In support of this idea, it was recently shown that an hourglass clock is unable to adjust dusk-peaking gene expression to peak at dusk in a summer condition and the misalignment of gene expression relative to the environment has negative consequences for survival to UV stress (6). Importantly, others have found that a selection can still occur between self-sustained clocks that establish similar phase angles of entrainment, but differ in amplitude and resetting ability, indicating that other features of the clock may also be under selective pressure (5).

We were surprised to find no advantage of the WT strain in the low-light equinox and winter conditions despite differences in phase angle of entrainment. We hypothesized that photoperiod-dependent differences in ROS accumulation underlie the photoperiod-specific outcomes of the competition experiments based on previous research. The cyanobacterial clock plays a critical role in activating genes of the OPPP at dusk to provide a non-photosynthetic source of NADPH required for clearance of ROS that accumulates across the day (14). As expected, shorter days, with fewer hours of photosynthesis, produced markedly lower ROS levels at dusk. Critically, by independently manipulating ROS through changing the light intensity or gas exchange, we were able to experimentally decouple ROS accumulation from photoperiod length. These manipulations revealed that ROS accumulation, rather than photoperiod, determined the fitness outcomes: cell survival and fitness in higher ROS environments, but not lower ROS environments, require an acrophase of dusk-peaking gene expression to occur close to dusk. Our findings highlight that the circadian clock plays a central role in helping cells cope with predictable environmental stress. Cyanobacteria maintained under constant laboratory conditions have, on occasion, lost circadian function, indicating that the clock imposes a physiological cost that can be outweighed by other selective pressures (18, 19). A central question in circadian biology is what aspects of the rhythmic environment impose a selection for circadian rhythmicity? Prior work showed that winter photoperiods promote lipid saturation and enhance survival to a cold shock and this response requires an intact circadian clock (7). Our results extend this principle by showing that the clock also protects cells against the daily accumulation of ROS, which reaches higher levels under summer photoperiods. Furthermore, the advantage conferred by the circadian system relies on tuning the FRP length. By phasing OPPP activity to coincide with dusk, when ROS burden peaks, the WT FRP positions ROS coping mechanisms precisely when they are most needed across all seasonal photoperiods. Together, these observations demonstrate that the WT FRP matches the phases of stress-response pathways to meet predictable environmental challenges.

## Materials and Methods

### Strains, plasmids and molecular methods

*E. coli* strains were grown at 37°C in lysogeny broth (LB, Lennox) in liquid culture or on agar, supplemented with the following antibiotics as needed: 20 μg/mL spectinomycin (Sp) plus 20 μg/mL streptomycin (Sm), 15 μg/mL gentamycin (Gm), 17 μg/mL chloramphenicol (Cm), 50 μg/mL kanamycin (Km), and 50 μg/mL nourseothricin (Nt). Unless noted otherwise, *S. elongatus* PCC 7942 and its derivative strains were grown in BG-11 medium (20) as liquid cultures or on agar plates (1.5% agar) at 30 °C under continuous illumination. Culture media for recombinant cyanobacterial strains were supplemented as needed with 2 μg/mL Sp plus 2 μg/mL Sm, 2 μg/mL Gm, 7.5 μg/mL Cm, 5 μg/mL Km, and 5 μg/mL Nt.

A complete list of plasmids and *S. elongatus* strains can be found in Tables S1 and S2. FRP strains carry a modified KaiC and an antibiotic resistance cassette in the *kaiC* native locus. Bioluminescent reporter strains contain the firefly luciferase gene downstream of either the P_*2336*_ or the P*kaiBC* promoter in neutral site NSII (21). The promoter sequences can be found in Table S7. Note that, while the P*kaiBC* construct includes luciferase fused in-frame to the first 27 base pairs of *kaiB*, the P_*2336*_ construct is not a translational fusion. Recombinant strains of *S. elongatus* were constructed using standard protocols (21). Briefly, fresh *S. elongatus* cultures were grown to an OD_750_ of 0.3-0.5 and 10 mL were pelleted by centrifugation at 5,000*xg*. The pelleted cells were washed with 1.5 mL of 10 mM NaCl and pelleted again. The cells were resuspended in 200 µL of BG-11 and incubated in the dark overnight with 200 ng of the recombinant plasmid. The cells were plated on BG-11 agar plates supplemented with the appropriate antibiotics and grown under continuous light until individual colonies appeared. Clones were tested for segregation using PCR. Unless noted otherwise, independent clones were used as biological replicates.

### Bioluminescence

Bioluminescence was monitored for several days (as indicated for each experiment) under the specified seasonal photoperiod followed by several days in constant light. The temperature was kept at 30°C and the light intensity was ∼15 µmol m^-2^ s^-1^. As previously described (9), the data were analyzed by fitting parabolas to the first and second peak in constant light for each biological replicate (Figure S2). Period was calculated as the difference in peak times, and phase relative to dusk was calculated by subtracting the time that dusk would have occurred on the first day in constant light from the first peak time.

Here, biological replicates were defined as independently grown samples from the same clone. Results similar to Figure 1D-F were obtained when additional clones were included (Figure S3).

### Growth-competition experiments

The relative fitness of the long-period and short-period strains were assessed by leveraging different antibiotic resistances of each strain. The long-period strain carries a Gm-resistance cassette and the short-period strain carries an Nt-resistance cassette. The axenic cultures were grown to an OD_750_ of 0.3-0.6 in constant light, washed three times with BG-11 via centrifugation at 5,000 × g for 10 min to remove antibiotics, and diluted with BG-11 to an OD750 of ∼0.1. The axenic cultures were then entrained to the specified seasonal photoperiod until they reached an OD of 0.3-0.6. Cultures were then mixed in a 1:1 ratio to a final OD_750_ of 0.1 and grown without antibiotics in the specified seasonally photoperiod for 8 generations. Every 2 generations, samples were taken to determine the ratio of each strain in the culture and the cultures were diluted to an OD_750_ of 0.1. To assay survival of each genotype, cultures were diluted using a 1:5 dilution series and plated in 4.5-μL spots in duplicate onto BG-11 agar with either Gm or Nt. Colony-forming units (CFU) of each strain were calculated across the two antibiotic regimes and were used to determine the percentage of each strain in the mixed population over time. To ensure that the expression of the antibiotic markers did not influence fitness, WT cultures that carried each drug-resistance marker adjacent to the *kaiABC* locus were grown together and assayed and displayed no differences in fitness (Figure S4). Additionally, we compared growth of the competition cultures on plates with and without antibiotics to test whether the strains grew equally well on both. We found that there was no significant difference between the CFU on the non-antibiotic plate compared to the sum of the CFU between both the Nt and Gm plates (student’s two sample paired t-test, p=0.47, Figure S5).

Unless otherwise specified, low light is defined as ∼40 µmol m^-2^ s^-1^ and high light as ∼70 µmol/m^2^/s. Strains were grown in 100 mL of media in 250 mL flasks with cotton plugs and were shaking at 120 rpm. For the summer-aerated condition, an aquarium pump with tubing was used to aerate the cultures.

### Quantification of ROS

ROS were quantified by using the fluorescent marker H_2_DCFDA (Thermo Fisher Scientific, no. D399), as previously described (14). Briefly, 2 mL of culture was collected and split into 1-mL aliquots. H_2_DCFDA stock solution was made by dissolving solid H_2_DCFDA in DMSO to a concentration of 1mM. The stock was added to one sample at a final concentration of 5 μM. Tubes were protected from light and shaken at 30 °C for 30 min. After incubation, 200 μL of each tube was added to a separate well in a 96-well plate. Fluorescence was quantified at an excitation of 480 nm and an emission of 520 nm on a Tecan Infinite M200 plate reader with the gain manually set to 120. Data were analyzed by subtracting untreated-sample background fluorescence from treated-sample fluorescence values and then normalizing by dividing by the OD_750_.

### Hydrogen Peroxide Stress

The survival ratio after exposure to hydrogen peroxide was assessed by adding hydrogen peroxide to a final concentration of 0.01% v/v to cells for 1 h starting 15 min before dusk and ending 45 min after dusk in a winter photoperiod. The concentration was chosen based on a pilot study that used 0.005% v/v, 0.01% v/v, and 0.025% v/v hydrogen peroxide. The concentration that produced a measurable response without resulting in 0% survival in any of the strains was 0.01% v/v. The cells were entrained to the winter photoperiod prior to the experiment and grown to an OD_750_ of ∼0.2-0.3. After the incubation time, cultures were diluted using a 1:5 dilution series and plated in 4.5-μL spots in duplicate onto BG-11 agar without antibiotic. Plates were left in constant illumination until colonies formed. Untreated cells were processed in parallel as controls. CFUs of each strain were calculated across the treated and untreated cultures and the survival ratio was calculated as the CFU on plates with treated cells divided by the CFU on plates with untreated cells. Biological replicates were defined as independently grown cultures utilizing three distinct clonal lineages.

## Data and Software Availability

All data are included in the article and/or SI Appendix

## Acknowledgments

A.N. is a San Diego IRACDA Scholar supported by the National Institutes of Health (NIH)/NIGMS K12 GM068524 Award. Research reported in this publication was supported by the National Institute of General Medical Sciences of the National Institutes of Health under Award Number R35GM118290. The content is solely the responsibility of the authors and does not necessarily represent the official views of the National Institutes of Health.

## References

1. T. Roenneberg, M. Merrow, Life before the clock: modeling circadian evolution. J Biol Rhythms 17, 495–505 (2002).

2. M. L. Jabbur, C. Dani, K. Spoelstra, A. N. Dodd, C. H. Johnson, Evaluating the Adaptive Fitness of Circadian Clocks and their Evolution. J Biol Rhythms 10.1177/07487304231219206, 7487304231219206 (2024).

3. Y. Ouyang, C. R. Andersson, T. Kondo, S. S. Golden, C. H. Johnson, Resonating circadian clocks enhance fitness in cyanobacteria. Proc Natl Acad Sci U S A 95, 8660–8664 (1998).

4. M. A. Woelfle, Y. Ouyang, K. Phanvijhitsiri, C. H. Johnson, The adaptive value of circadian clocks: an experimental assessment in cyanobacteria. Curr Biol 14, 1481–1486 (2004).

5. C. Dani et al., Seasonal cycles select for self-sustained circadian oscillators. Curr Biol 10.1016/j.cub.2026.01.012 (2026).

6. A. F. Schober, C. Holmes, S. Palmer, M. J. Rust, The free-running property of circadian clocks is needed to tolerate changing photoperiods. Curr Biol 10.1016/j.cub.2026.01.066 (2026).

7. M. L. Jabbur, B. P. Bratton, C. H. Johnson, Bacteria can anticipate the seasons: Photoperiodism in cyanobacteria. Science 385, 1105–1111 (2024).

8. M. Fang, C. L. Partch, A. LiWang, S. S. Golden, Prokaryotic Circadian Systems: Cyanobacteria and Beyond. Annu Rev Microbiol 79, 523–545 (2025).

9. E. Leypunskiy et al., The cyanobacterial circadian clock follows midday in vivo and in vitro. Elife 6 (2017).

10. K. Greenham et al., Geographic Variation of Plant Circadian Clock Function in Natural and Agricultural Settings. J Biol Rhythms 32, 26–34 (2017).

11. M. A. Hardigan et al., Genome diversity of tuber-bearing Solanum uncovers complex evolutionary history and targets of domestication in the cultivated potato. Proc Natl Acad Sci U S A 114, E9999–E10008 (2017).

12. C. S. Pittendrigh, T. Takamura, Latitudinal clines in the properties of a circadian pacemaker. J Biol Rhythms 4, 217–235 (1989).

13. M. Ishiura et al., Expression of a gene cluster kaiABC as a circadian feedback process in cyanobacteria. Science 281, 1519–1523 (1998).

14. S. Diamond et al., Redox crisis underlies conditional light-dark lethality in cyanobacterial mutants that lack the circadian regulator, RpaA. Proc Natl Acad Sci U S A 114, E580–E589 (2017).

15. H. Min, S. S. Golden, A new circadian class 2 gene, opcA, whose product is important for reductant production at night in Synechococcus elongatus PCC 7942. J Bacteriol 182, 6214–6221 (2000).

16. J. R. Piechura, K. Amarnath, E. K. O’Shea, Natural changes in light interact with circadian regulation at promoters to control gene expression in cyanobacteria. Elife 6 (2017).

17. J. S. Markson, J. R. Piechura, A. M. Puszynska, E. K. O’Shea, Circadian control of global gene expression by the cyanobacterial master regulator RpaA. Cell 155, 1396–1408 (2013).

18. M. Adomako et al., Comparative Genomics of Synechococcus elongatus Explains the Phenotypic Diversity of the Strains. mBio 13, e0086222 (2022).

19. A. Mendana et al., Mutations in the circadian cycle drive adaptive plasticity in cyanobacteria. Proc Natl Acad Sci U S A 122, e2506928122 (2025).

20. R. Rippka, Isolation and purification of cyanobacteria. Methods Enzymol 167, 3–27 (1988).

21. E. M. Clerico, J. L. Ditty, S. S. Golden, Specialized techniques for site-directed mutagenesis in cyanobacteria. Methods Mol Biol 362, 155–171 (2007).

