## Supplemental Figures and Tables for "Circadian phase alignment promotes survival under predictive oxidative stress"

**This PDF file includes:**

Figures S1 to S5
Tables S1 to S6
SI References

### Figures

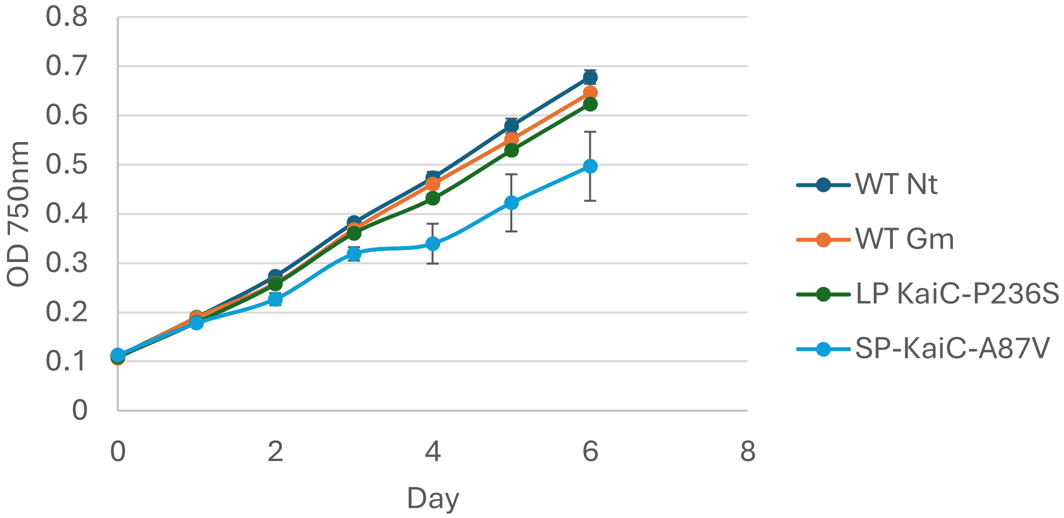


**Fig. S1.** Growth curves for all strains in a summer low-light condition. Data represent mean±SEM, n=3 for all data points.

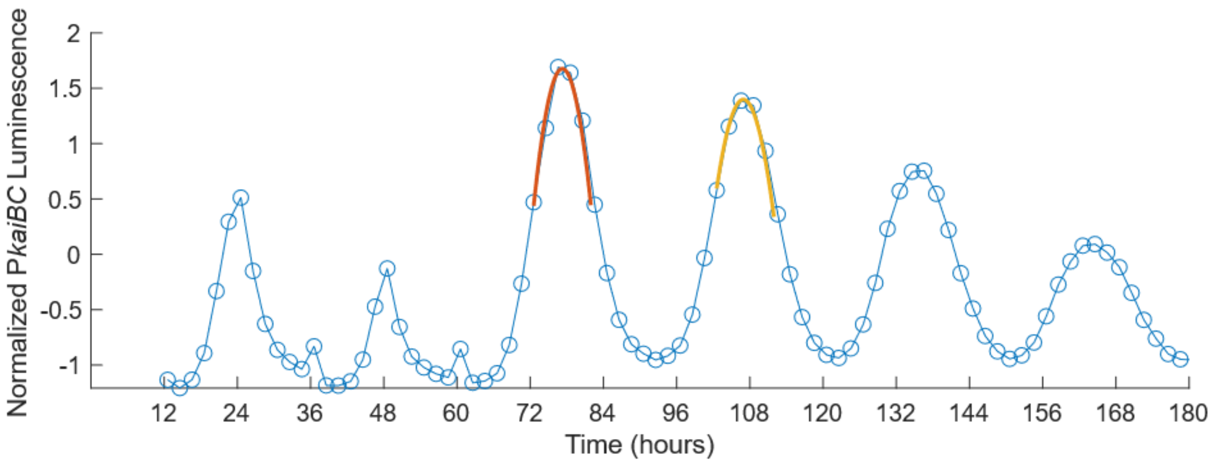


**Fig. S2.** Representative plot of parabolic fitting. Parabolas were fit to the first (red) and second (yellow) peaks in constant light for each biological replicate. Period was calculated as the difference in peak times and phase was calculated by subtracting the time that dusk would have occurred on the first day in constant light from the first peak time.

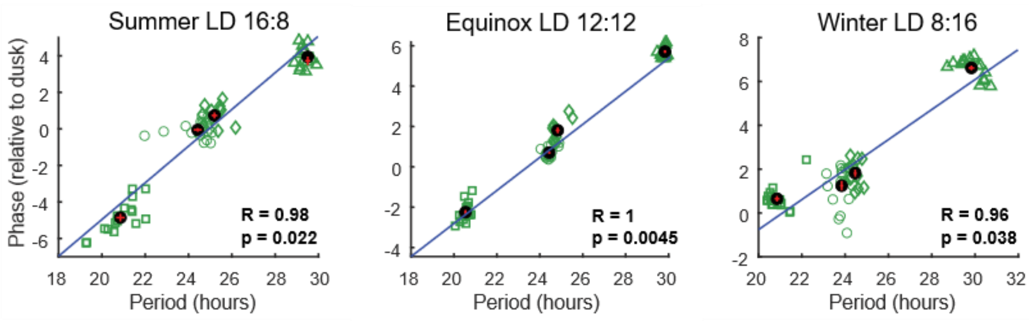


**Fig. S3.** Additional clones were tested to show the FRP is associated with phase of expression from P*kaiBC* in all seasonal photoperiods. Pearson's correlation was used to calculate R and p-values for linear correlation. The green symbols represent individual replicates (○ WTNt, ◊ WTGm, □ SP KaiC-A87V, Δ LP KaiC-P236S). The black dots represent the mean of each strain. The red lines represent the SEM, For summer and winter: $n_{SP}$=17, $n_{LP}$=17, $n_{WTGm}$=18, $n_{WTNt}$=12. For equinox: $n_{SP}$=14, $n_{LP}$=14, $n_{WTGm}$=14, $n_{WTNt}$=11.

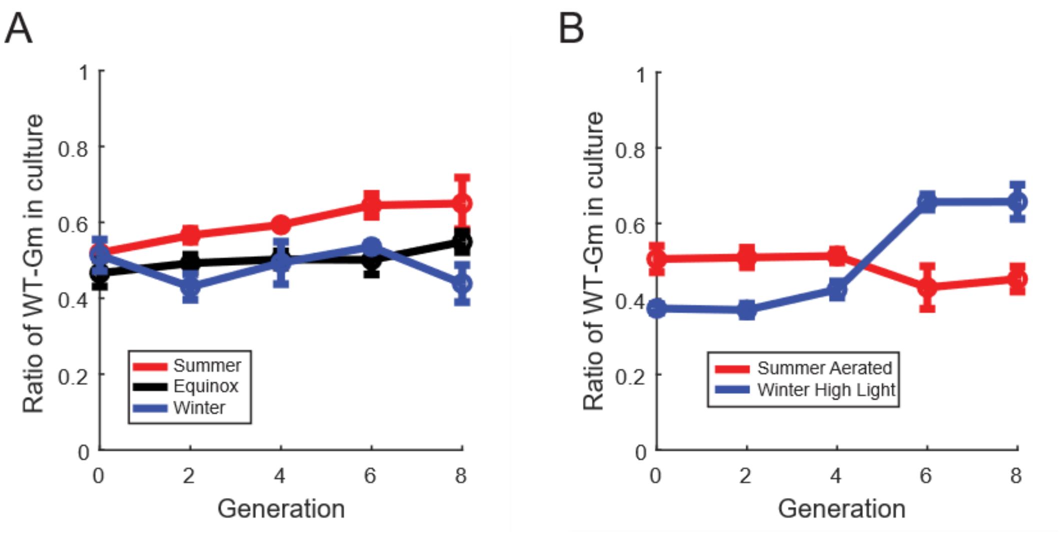


**Fig. S4.** Growth competition experiments between WT strains with either Nt or Gm antibiotic resistances. Results of the competition in conditions matching that of A.) Figure 1 and B.) Figure 3E, F. Data represent mean±SEM, n=3 for all datapoints.

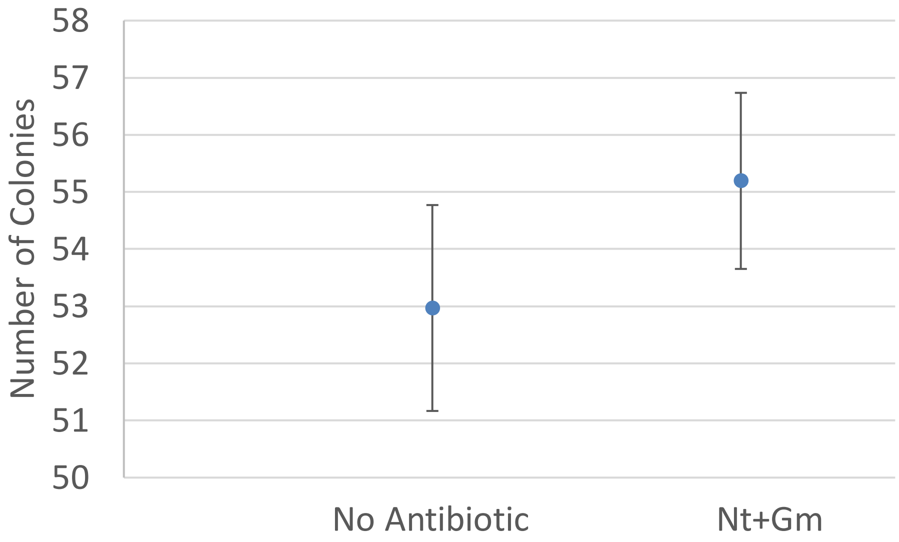


**Fig. S5.** There is no significant difference between the number of colonies that grow on no antibiotic plates and the sum of those that grew on the Nt and Gm plates (student's two-sample paired t-test, $p=0.47$). Data represent mean$\pm$SEM, $n=16$ for all datapoints.

**Tables**

**Table S1.** Plasmids used in this study

| Plasmid | Description | Antibiotic | Source |
|---|---|---|---|
| pCkaiABC | *KaiABC*+$SpSm^R$ flanked by homologous arms to recombine at native locus | SpSm | Ishiura *et al.*, 1998 (1) |
| pAM5979 | Modified pCKaiABC to provide Gm resistance; *KaiABC*+$Gm^R$ flanked by homologous arms to recombine at native locus | Gm | Lab collection |
| pAM5980 | Modified pCKaiABC to provide Nt resistance; *KaiABC*+$Nt^R$ flanked by homologous arms to recombine at native locus | Nt | Lab collection |
| pAM5981 | pAM5979 modified to make the long-period mutant KaiC T409A | Gm | Lab collection |
| pAM5982 | pAM5980 modified to make the short-period mutant KaiC R321Q | Nt | Lab collection |
| pAM5994 | pAM5979 modified to make the long-period mutant KaiC P236S | Gm | Lab collection |
| pAM5995 | pAM5980 modified to make the short-period mutant KaiC A87V | Nt | Lab collection |
| pAM5999 | pAM5979 modified to make the short-period mutant KaiC A87V with Gm resistance | Gm | Lab collection |
| pAM6015 | $P_{2336}$-*luciferase* with Km resistance flanked by NSII homologous arms | Km | Lab collection |

**Table S2.** Cyanobacterial strains used in this study

| Strain | Description | Antibiotic | Source |
|---|---|---|---|
| AMC1751 | WT AMC541 strain ($P_{kaiBC}$-*luciferase* in NSII) transformed with pAM4252 to inactivate KaiABC | Km, Cm | Lab collection |
| AMC2798 | AMC1751 transformed with pAM5979; WT period strain with $Gm^R$ | Gm, Cm | Lab collection |
| AMC2799 | AMC1751 transformed with pAM5980; WT period strain with $Nt^R$ | Nt, Cm | Lab collection |
| AMC2806 | AMC1751 transformed with pAM5994; Long-period KaiC P236S strain with $Gm^R$ | Gm, Cm | Lab collection |
| AMC2807 | AMC1751 transformed with pAM5995. Short-period KaiC A87V strain with $Nt^R$ | Nt, Cm | Lab collection |
| AMC2834 | AMC2798 transformed with pAM6015 to replace $P_{kaiBC}$-*luciferase* in NSII with $P_{2336}$-*luciferase* | Gm, Km | Lab collection |
| AMC2835 | AMC2799 transformed with pAM6015 to replace $P_{kaiBC}$-*luciferase* in NSII with $P_{2336}$-*luciferase* | Nt, Km | Lab collection |
| AMC2838 | AMC2806 transformed with pAM6015 to replace $P_{kaiBC}$-*luciferase* in NSII with $P_{2336}$-*luciferase* | Gm, Km | Lab collection |
| AMC2839 | AMC2807 transformed with pAM6015 to replace $P_{kaiBC}$-*luciferase* in NSII with $P_{2336}$-*luciferase* | Nt, Km | Lab collection |

**Table S3.** Two Way ANOVA results for ROS accumulation at dusk

| Source | Sum Sq. | d.f. | Mean Sq. | F | Prob>F |
|---|---|---|---|---|---|
| Strain | 5.65E+07 | 3 | 1.88E+07 | 1.41 | 0.26 |
| Condition | 3.16E+09 | 3 | 1.05E+09 | 78.65 | 7.45E-15 |
| Strain:Condition | 2.20E+08 | 9 | 2.45E+07 | 1.83 | 0.10 |
| Error | 4.29E+08 | 32 | 1.34E+07 | | |
| Total | 3.87E+09 | 47 | | | |

**Table S4.** Tukey's posthoc test of condition following Two-Way ANOVA of ROS accumulation at Dusk

| Group 1 | Group 2 | Estimated mean difference | CI Lower bound | CI upper bound | p-value |
|---|---|---|---|---|---|
| Summer | Summer Aerated | 1.75E+04 | 2.15E+04 | 2.56E+04 | 8.42E-15 |
| Winter | Summer | -2.05E+04 | -1.65E+04 | -1.24E+04 | 1.18E-11 |
| Winter High Light | Summer | -1.31E+04 | -9.05E+03 | -5.00E+03 | 5.42E-06 |
| Winter | Summer Aerated | 1.02E+03 | 5.07E+03 | 9.12E+03 | 9.56E-03 |
| Winter High Light | Summer Aerated | 8.45E+03 | 1.25E+04 | 1.66E+04 | 8.77E-09 |
| Winter | Winter High Light | -1.15E+04 | -7.43E+03 | -3.38E+03 | 1.23E-04 |

**Table S5.** One-way ANOVA results for survival after exposure to hydrogen peroxide at dusk

| Source | Sum of Squares | df | Mean Squares | F | p-value |
|---|---|---|---|---|---|
| Groups | 1.07 | 3 | 0.36 | 4.44 | 0.011 |
| Error | 2.26 | 28 | 0.081 | | |
| Total | 3.33 | 31 | | | |

**Table S6.** Tukey's posthoc test following One-Way ANOVA of survival after exposure to hydrogen peroxide at dusk

| Group 1 | Group 2 | Lower CI | Estimate | Upper CI | p-value |
|---|---|---|---|---|---|
| WT Nt | WT Gm | -0.34 | 0.050 | 0.44 | 0.99 |
| WT Nt | LP | 0.062 | 0.45 | 0.84 | 0.018 |
| WT Nt | SP | -0.090 | 0.30 | 0.68 | 0.18 |
| WT Gm | LP | 0.012 | 0.40 | 0.79 | 0.041 |
| WT Gm | SP | -0.14 | 0.25 | 0.64 | 0.32 |
| LP | SP | -0.54 | -0.15 | 0.24 | 0.71 |

**Table S7.** Promoter sequences for bioluminescence monitoring

| Promoter | Sequence 5'→ 3' |
|---|---|
| *kaiBC* (including the first 27 base pairs of *kaiB*) | AAAGGTAAAGGAGGTCTTAAGCTCGGCTCAATTTCTCTCTTTATCCTGTT<br>AGATGGTTTGATTGCTGTTGCTACCCCGTTGATCTGCGTATGAGCCCTC<br>GTAAAACCTACATTCCC |
| *2336* | GAGGGGCGATCGCCGAGGGCCAGAGCGTCAACGTTAGCCTGCGAAAA<br>GATTGGCCCAGCCTTTGTTTTGCTGAATGTTCCCGCCATTCGTTAAATCA<br>GTTGGTGAGATTTGAGATCCCTGACAGAGCGATCGCTAATCTCTGAACTA<br>AATGCTCATGCTTGCTTAGCTTTCCCCCAAGCCTTCACCTCACTTGTACAG<br>GAGGGGTAT |

**SI References**


1. M. Ishiura *et al.*, Expression of a gene cluster kaiABC as a circadian feedback process in cyanobacteria. *Science* **281**, 1519-1523 (1998).
